# Molecular insights into human Shieldin complex assembly and recruitment to DSBs

**DOI:** 10.1101/2022.06.09.495453

**Authors:** Vivek Susvirkar, Alex C. Faesen

## Abstract

The Shieldin complex represses end resection at DNA double-strand breaks (DSBs) and thereby serves as a pro-non homologous end joining (NHEJ) factor in the G1 phase of the cell cycle. Its components SHLD1, SHLD2, SHLD3 and REV7 are recruited in a hierarchical fashion. SHLD3 and REV7 localize first to DSBs, while the subsequently recruited SHLD2 is the only known DNA binding protein in the complex. The molecular details of the initial recruitment of SHLD3 and REV7, and the subsequent assembly of Shieldin on DSBs are unclear. Here, we report the identification of a promiscuous DNA binding domain in the C-terminal half of SHLD3. At the N-terminus, SHLD3 interacts with a dimer of REV7 molecules. We show that the interaction between SHLD3 and the first REV7 is remarkably slow, which is likely due to the substantial activation energy required to remodel mobile structural elements within the REV7 molecule to allow for binding to SHLD3. In contrast, the interaction between SHLD3 and SHLD2 with a second REV7 molecule is fast and does not require structural remodelling. Overall, these results provide insights into the rate-limiting step of the molecular assembly and recruitment of Shieldin complex at DNA DSBs.

## Introduction

DNA double-strand breaks (DSBs) are highly toxic to cells as they cause full rupture of the chromosomes. Incorrect or a lack of repair leads to genomic anomalies ranging from insertion, deletions, duplications and translocations. These anomalies have been linked to embryonic death, early aging, genetic disorders, immunodeficiency, neurological disorders and cancer. During the G1 phase of cell cycle, the Shieldin complex is recruited in a 53BP1-RIF1 dependent manner and binds ssDNA to shield it from DNA resection (Dev et al., 2018; Ghezraoui et al., 2018; Gupta et al., 2018; Noordermeer et al., 2018). This commits the repair of DNA to the non-homologous end joining (NHEJ) pathway and thereby guards genomic integrity in an event where homology repair (HR) would be detrimental. The Shieldin complex has been identified as a key factor responsible for PARPi resistance in tumour cells lacking 53BP1. Therefore, understanding molecular mechanism of Shieldin complex-mediated repair by NHEJ of DSBs is essential to provide better insights in the regulation of DNA repair in health and disease.

The Shieldin complex consists of four subunits, SHLD1 (205 residues), SHLD2 (835 residues), SHLD3 (250 residues), and HORMA domain REV7 (211 residues, also known as MAD2L2). The SHLD3-REV7 module is considered to be the recruitment arm, while SHLD2-SHLD1 is the effector arm of the complex (Setiaputra and Durocher, 2019). The assembly of the Shieldin complex in cells follows a linear hierarchy where SHLD3 localizes first to DNA breaks. This is followed by REV7 and subsequently by SHLD2-SHLD1 in a 53BP1 and RIF1-dependent manner (Dev et al., 2018; Gupta et al., 2018). SHLD2 is proposed to function as a scaffold protein and contains three putative oligonucleotide-binding (OB) folds which have been shown to bind ssDNA (Dev et al., 2018; Noordermeer et al., 2018). The functions of SHLD1 and SHLD3 are unclear as they are biochemically uncharacterized. The molecular details of how Shieldin complex assembly is regulated and how it recognizes DSBs in order to protect them from end resection remain incomplete.

REV7 is a member of the HORMA (HOP1, REV7, MAD2) domain family, which are conserved signalling proteins serving as sensors in a variety of pathways, ranging from bacterial immunity, eukaryotic cell cycle, genome stability, sexual reproduction, and cellular homeostasis (Gu et al., 2022). HORMA domain proteins regulate the assembly of effector complexes by reversibly changing its protein’s three-dimensional structure to control its protein-protein interaction potential. Conversion between these conformer states in REV7 (‘open’ and ‘closed’) is spontaneous but slow, likely due to the substantial unfolding and refolding of mobile elements to the core fold of the HORMA domain (Clairmont et al., 2020). Factors that regulate or catalyse the conversion between conformer states, directly control the rate-limiting step in the assembly and disassembly of these effector complexes at the right place and time (Faesen et al., 2017; Gu et al., 2022; Musacchio, 2015)

Central to the unusual interaction mechanism of REV7 is the extraordinary ability to wrap its C-terminal tail (‘seat-belt’) around an interacting peptide motif of a client protein, thereby creating a very stable complex. Both Polζ and Shieldin incorporate a dimer of REV7 molecules, which are essential for their function *in vitro* and in cells (de Krijger et al., 2021; Malik et al., 2020; Rizzo et al., 2018). Using the seat-belt mechanism, REV7 captures the REV7-binding-motifs (RBM) in SHLD3 (residues 49 to 62) and REV3 (RBM1, residues 1875 to 1896, and RBM2, residues 1991 to 2012) (Dai et al., 2020; Hara et al., 2010; Rizzo et al., 2018). Additionally, REV7 interacts with the REV7 interacting motif (RIM) present on SHLD2, which mediates the interaction between SHLD3 and a second REV7 molecule (Gupta et al., 2018; Liang et al., 2020). Dimerization is a common feature of HORMA domains. Asymmetric dimerization of an ‘open’ and ‘closed’ form of HORMA domain protein MAD2 was shown to be essential in the mechanism to catalyse the conversion between conformer states (Antoni et al., 2005; Faesen et al., 2017). After conversion, the MAD2 dimer dissociates, and the converted MAD2 molecule is incorporated in its effector complex (the Mitotic Checkpoint Complex). It is currently unclear if the stable incorporation of a dimer of REV7 molecules affects the assembly kinetics of Polζ and Shieldin.

Here, we study the molecular mechanism of Shieldin recruitment and assembly at DNA using a reconstituted recombinant human Shieldin complex consisting of full-length SHLD3, REV7 and the N-terminal peptide (1-95) of SHLD2. We show that this complex, which lacks the SHLD2 DNA binding domain, is able to bind both double- and single-strand DNA. We identify a DNA binding domain in the C-terminal part of SHLD3. Point mutants of conserved residues in a predicted electropositive pocket prevent DNA binding. Quantification of the interaction kinetics of REV7 with the RBM’s on SHLD3 and REV3, shows a conserved and extremely slow capture by the seat-belt of the HORMA domain. In contrast, the interaction of the second REV7 molecule to REV7-SHLD3-SHLD2 is fast. These observations support a model where the DNA binding domain contributes to the initial recruitment of SHLD3 and that the subsequent assembly of the Shieldin complex on DSBs is rate-limited by the incorporation of only the first REV7 molecule.

## Results

### SHLD3 has a DNA binding domain

To understand the molecular mechanism behind the recruitment of the Shieldin complex to sites of DSBs, we purified a stable ternary complex (called Shieldin hereafter) consisting of human full-length SHLD3 along with a N-terminal peptide containing the first 95 residues of SHLD2 (SHLD2^1-95^) and REV7 (Figure 1a). Although this complex lacks the DNA binding OB-fold domains in SHLD2, we noticed that the complex could bind to a heparin column, which suggests an ability to bind DNA (Figure 1b). To test for DNA binding, we incubated the Shieldin complex with an excess of 5(6)-carboxyfluorescein (5(6)-FAM) labelled single-strand DNA (ssDNA) and carried out analytical size exclusion chromatography (SEC). In the presence of the Shieldin complex, the elution peak of labelled ssDNA moved to a higher apparent molecular weight and co-migrated with Shieldin (Figure 1c). Next, we used the FAM labelled DNA to carry out fluorescence anisotropy (FA) measurements to quantify the binding affinity of the Shieldin complex for different DNA substrates. No major differences in dissociation constant (K_D_) were observed between ssDNA telomeric (200nM), ssDNA non-telomeric (137 nM), dsDNA telomeric (166 nM) and dsDNA non-telomeric (180 nM) (Figure 1d,e).

**Figure 1.**
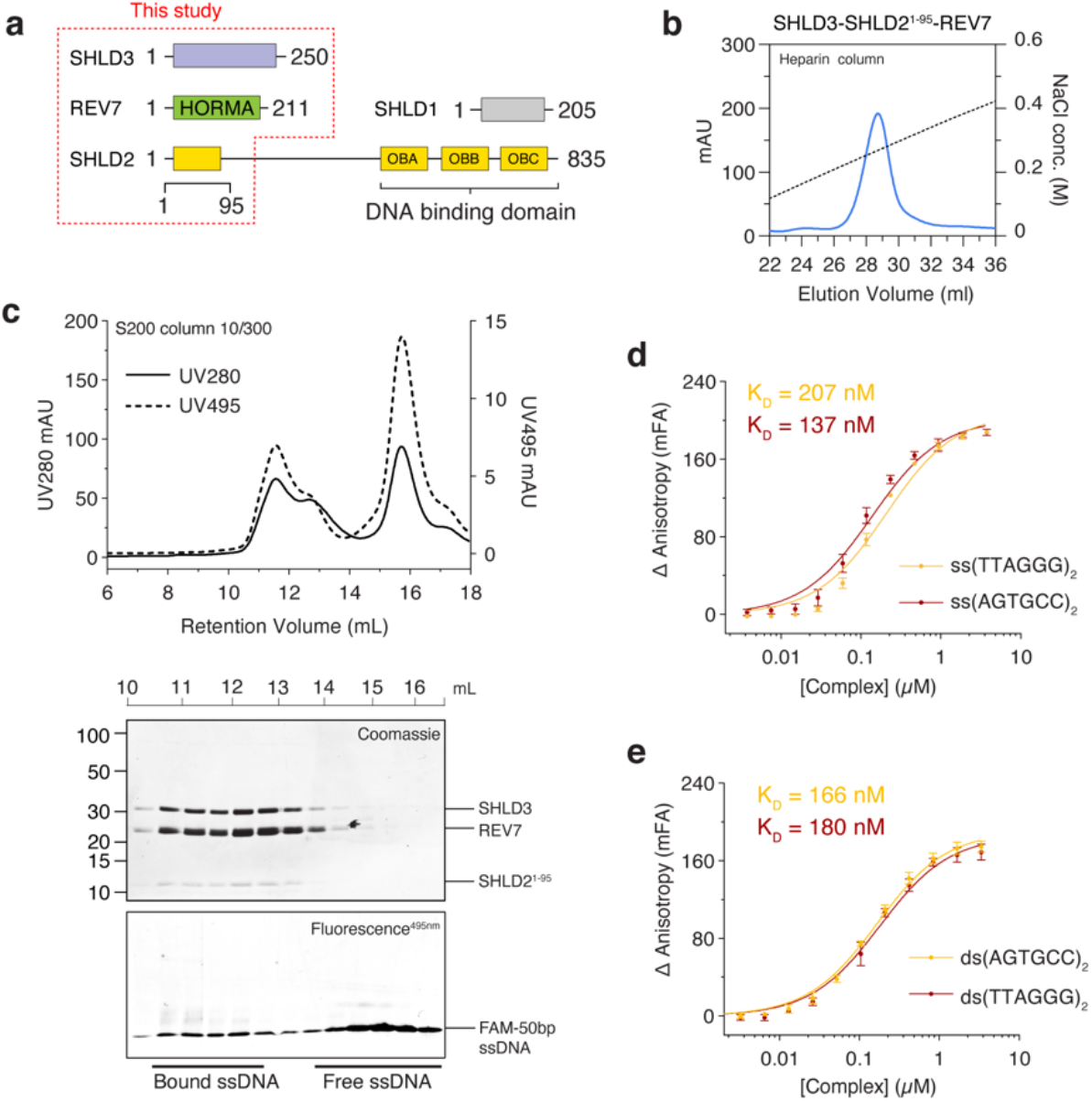
SHLD3-SHLD2^1-95^-REV7 ternary complex binds DNA. **a)** Schematic drawing of SHLD1, SHLD2, SHLD3 and REV7 proteins of the Shieldin complex. Protein constructs present in this study are highlighted. **b)** Heparin column elution profile of SHLD3-SHLD2^1-95^-REV7 ternary complex. **c)** Co-migration of ssDNA and SHLD3-SHLD2^1-95^-REV7 complex by size exclusion chromatography. **d)** Fluorescence anisotropy measurement of Shieldin complex for binding affinities against non-telomeric and telomeric ssDNA and **e)** for non-telomeric and telomeric dsDNA. Error bars represent s.d. (n = 3 independent experiments).

To determine which component in the Shieldin complex harbours this promiscuous DNA binding domain, we tested different subcomplexes of the Shieldin complex for DNA binding. After incubation with complexes lacking SHLD3, we observed that the ability to bind to DNA was lost (Figure S1a). Since the SHLD3 N-terminus is involved in REV7 binding we suspected that the DNA binding domain was C-terminal of the REV7 binding motif (Dai et al., 2020). To test this, we generated a truncation mutant that lacks this SHLD3 region (84-250) (Figure S1b). As expected, this Shieldin complex failed to bind ssDNA (Figure 2a). To understand the molecular mechanism of DNA recognition by SHLD3, we sought to identify and purify the DNA binding domain. Analysis of sequence conservation reveals high levels of conservation in the C-terminal region of SHLD3 (Ashkenazy et al., 2016). This coincided with predicted order in this region, suggesting a folded domain (Figure 2b) (Erdős et al., 2021). We carried out limited proteolysis with trypsin and observed two stable fragments of SHLD3 corresponding to ~15 kDa and ~13 kDa, which agrees with the expected mass of SHLD3^140-250^ (SHLD3^CTD^; Figure 2c). This SHLD3^CTD^ was expressed in *E.coli* and purified to homogeneity (Figure 2d). We tested it for DNA binding using a FA competition assay, where we pre-incubate SHLD3^CTD^ with labelled ssDNA and subsequently monitor the loss of FA signal in the presence of an excess of dark unlabelled competing ssDNA or dsDNA. This showed that indeed SHLD3^CTD^ can bind DNA with no discernible preference for ss- or dsDNA (Figure 2e). This was confirmed in titration experiments, where the measured binding constants of the tested DNA constructs were similar to the Shieldin complex (Figure S1e,f). Next, we used Alphafold2 to model the structure of SHLD3^CTD^ (Jumper et al., 2021). SHLD3^CTD^ was modelled using ColabFold with default settings, which generated a high-confidence 3D model (pLDDT score of greater than 90) (Figure S1c) (Mirdita et al., 2021). The predicted model of SHLD3^CTD^ showed a high degree of homology to translation initiation factor EIF4-E, a known nucleotide binding protein (Figure S1d). Surface analysis showed the presence of an electropositive patch that includes the conserved residues H242 and K243 (Figure 2f-h). As expected, mutating H242 and K243 to alanine reduced DNA binding affinity for all substrates (Figure 2i and S1g). Taken together, we identify SHLD3 as a novel promiscuous DNA binding protein within the Shieldin complex.

**Figure 2.**
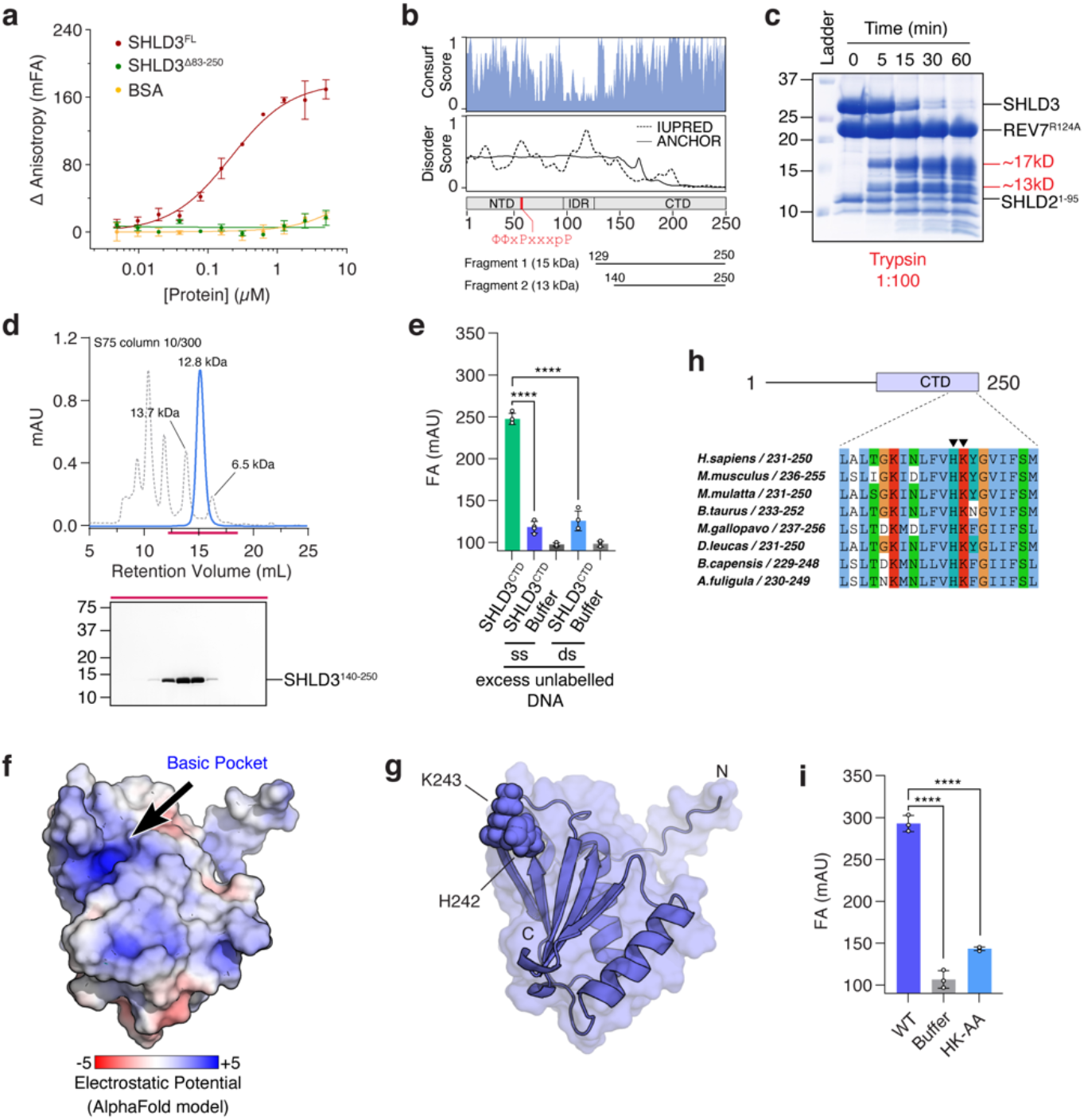
SHLD3 C-terminus contains a DNA binding domain. **a)** Fluorescence anisotrophy measurement of different constructs of Shieldin complex for binding affinities against ssDNA. Shieldin complex with deletion of SHLD3 residues 83-250 is deficient in DNA binding. Brown line represents Shieldin complex, green line represents Shieldin complex^ΔSHLD3(83-250)^, yellow line represents BSA. Error bars represent s.d. (n = 3 independent experiments). **b)** Disorder prediction of SHLD3 using IUPRED shows the N-terminal region (1-150) is disordered while the C-terminal region is ordered. The conservation score shows increased conservation in the N-terminal region (1-83) and C-terminal region (140-250). **c)** Proteolytic digestion of the Shieldin complex with trypsin shows degradation of SHLD3 to 15 kDa and 13 kDa fragments. **d)** Purification of SHLD3^140-250^ by size exclusion chromatography. **e)** FA competition assay shows SHLD3^CTD^ binds ssDNA and dsDNA with no preference.. FAM-labelled ssDNA at 10 nM was incubated with 1 μM SHLD3^CTD^. Labelled ssDNA was competed out using either 1 μM non-telomeric ssDNA or dsDNA. Error bars represent s.d. (n = 4 independent experiments). Two-tailed Student’s test are indicated: ****p<0.0001. **f)** Surface electrostatic analysis of SHLD3^CTD^ AlphaFold2 model reveals presence of an electropositive patch. **g)** SHLD3^CTD^ shown in surface representation with H242 and K243 shown as spheres. **h)** Residues H242 and K243 are well conserved across SHLD3 homologues in higher eukaryotes. Sequence alignment was performed with Clustal package in Jalview (Sievers et al., 2011; Waterhouse et al., 2009). Residues are coloured according to Clustalx scheme, where conserved residues are coloured as follows: blue (hydrophobic), red (positively charged), orange (glycine), cyan (hydrophobic) and green (polar). **i)** Introduction of H242A and K243A point mutants abolishes binding affinity of SHLD3^CTD^ to ssDNA.

### The Shieldin complex contains an asymmetric REV7 dimer

The Shieldin complex stably integrates a REV7 dimer (Liang et al., 2020). Using SEC coupled to static angle light scattering equipment (SEC-SLS), we measure a 90 kDa mass for the Shieldin complex, which agrees with a theoretical mass of 88 kDa of a 2:1:1 stoichiometric REV7-SHLD3-SHLD2^1-95^ complex (Figure S2a). Following a path previously explored with MAD2, we created REV7 mutants with the aim to trap them in a single conformer and subsequently used a surface-charge based anion exchange assay to identify and purify separate conformers (Mapelli et al., 2007) (Figure 3a and S2b). Elution from the anion exchange column correlates with the conformer identity (‘open’ or ‘closed’) (Clairmont et al., 2020; Mapelli et al., 2007). All of the tested REV7 constructs eluted as distinct conformers, and only the REV7 ‘loopless’ (REV7^LL^) eluted as an ‘open’ REV7 (Figure 3b). We did not observe conformer mixtures or changes in conformer identity after incubations over prolonged times at different temperatures (Clairmont et al., 2020; Luo et al., 2004; Mapelli et al., 2007). We therefore concluded that these are thermodynamically the most stable conformers in the unbound state.

**Figure 3.**
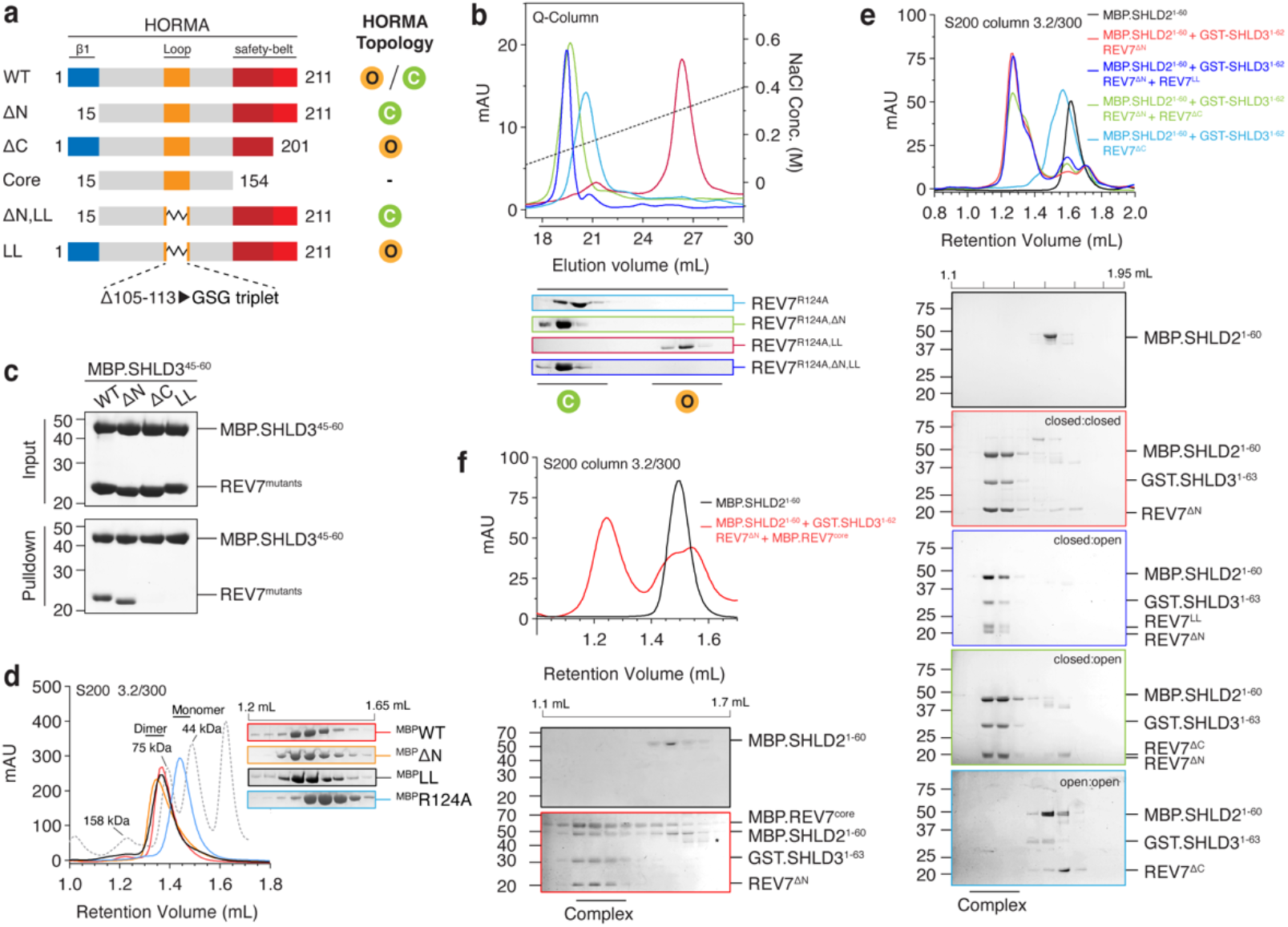
Shieldin complex contains REV7 asymmetric dimer. **a)** Schematic drawing of REV7 constructs used in this study to trap REV7 in different topologies. **b)** Anion exchange chromatography separates monomeric ‘closed’ REV7 from ‘open’ REV7. REV7, REV7^ΔN^, REV7^ΔN,LL^ elute in the ‘closed’ conformation. REV7^LL^ elute in the ‘open’ REV7 conformation. The R124A mutant was used to eliminate contributions of dimerization. Eluted fractions were collected and analysed by SDS-PAGE. **c)** MBP pulldown showing safety-belt interaction between SHLD3^45-65^ with ‘closed’ REV7 mutants. ‘Open’ REV7 mutants fail to bind SHLD3^45-65^ in safety belt conformation. **d)** Elution profile of MBP-REV7 protein mutants from size exclusion chromatography. The consecutive 50 μL fractions eluting from 1.2 and 1.65 mL are shown. **e)** Size exclusion chromatography profiles show interaction between equimolar ratios of MBP-SHLD2^1-60^, REV7^ΔN/ΔC^, GST-SHLD3^1-62^, and REV7^ΔN/ΔC/LL^. Samples were analysed by SDS-PAGE. **f)** Size exclusion chromatography profiles show interaction between MBP-SHLD2^1-60^, REV7^ΔN^, GST-SHLD3^1-62^ and REV7^core^. MBP-SHLD2^1-60^ and REV7^core^ were loaded in excess. Samples were analysed by SDS-PAGE.

Next, we tested whether the mutants could bind the RBM of SHLD3. As expected, the REV7^WT^ and REV7^ΔN^ constructs, which both eluted as ‘closed’ REV7, could interact with SHLD3^45-60^ (Figure 3c). Conversely, REV7^LL^ and REV7^ΔC^, which are trapped in the ‘open’ state, could not (Clairmont et al., 2020). This confirms that REV7 requires both a functional seatbelt and the ability for this seatbelt to adopt the closed conformation to interact to the RBM on SHLD3 (Clairmont et al., 2020; Dai et al., 2020). We subsequently investigated whether REV7 dimerization is affected by the conformer mutants. Since REV7^WT^ weakly homodimerizes with a K_D_ of ~ 2 μM, we injected our conformer mutants on the SEC column using a concentration of 20 μM (Rizzo et al., 2018). All REV7 conformer constructs were able to dimerize, which contrasts the conformer sensitive dimerization observed for MAD2 (Mapelli et al., 2007; Yang et al., 2008) (Figure 3d and S2c).

Having established that the first REV7 molecule has to be in the closed state, we wondered if the incorporation of the second REV7 in Shieldin is also conformer-sensitive. To test this, we monitored assembly of Shieldin by mixing purified individual components in stoichiometric amounts. We used MBP tagged SHLD2^1-60^ to circumvent stability issues with the full-length protein. As expected, when using mutants that can only be in the ’open’ conformer, we could not observe assembly of the Shieldin complex, confirming that at least one REV7 molecule needs a functional seatbelt in the closed conformation (Figure 3e). In contrast, complex formation was unperturbed when using REV7^ΔN^ (which forms ‘closed’:’closed’ dimers), REV7 ^ΔN^-REV7^LL^ or REV7^δN^-REV7^δC^ (which both form ‘closed’:’open’ dimers). This shows that there is no discrimination between conformer states for the second REV7 molecule, suggesting that the mobile elements of the second REV7 are not essential. To test this, we created a REV7^core^ construct that lacks all mobile elements and therefore cannot adopt the ‘closed’ or the ‘open’ conformer (Figure 3a and S2b). As expected, this REV7^core^ construct failed to interact with SHLD3^RBM^ (Figure S2d). In contrast, when incubated with MBP-SHLD2 and a preformed REV7-SHLD3, the REV7^core^ can be incorporated in Shieldin as judged by the shift of the SHLD2-elution peak and co-migration with the other components (Figure 3f). Taken together, we show that the Shieldin complex contains an asymmetric REV7 dimer, with one ‘closed’ REV7 bound to SHLD3 and a second previously unidentified state of REV7 distinct from the canonical ‘open’ and ‘closed’ states where the interaction with SHLD2 is mediated via the static core of REV7.

### Only REV7-SHLD3^RBM^ binding is rate-limiting for Shieldin assembly

The emerging paradigm for HORMA domain proteins is that they default to an inactive ‘open’ state, which is than poised to convert to an ‘closed’ partner-bound active state. This slow, but spontaneous, conversion is a rate-limiting step in the assembly of the respective effector complex. Indeed, we previously showed that REV7-SHLD3 complex formation takes several hours (de Krijger et al., 2021). However, as described in Figure 3b, the purified REV7 is already in the ‘closed’ state, so potentially no conversion would be necessary. Indeed, a recent study on MAD2 describes the possibility of threading a short flexible client peptide, like the RBM on SHLD3, into a pre-closed HORMA domain (Piano et al., 2021). Since this threading was about 2-orders of magnitude faster than the canonical capture after converting from the ‘open’ state to the ‘closed’ state, we wondered how the interaction kinetics of a pre-closed apo-REV7 to SHLD3^RBM^ compared to known HORMA domain interaction kinetics.

To quantify the binding rates, we used surface plasmon resonance (SPR) with immobilized SHLD3^1-62^ (Figure 4a-c) or REV3^1871-2014^ (Figure 4d,e and S3b) and injected either wild-type or dimerization incompetent REV7 (REV7^R124A^). This showed that REV7^WT^ interacts slowly to SHLD3^1-62^. When washing with buffer to measure the dissociation of REV7, only a fraction of the signal is lost, suggesting only a partial disassembly of the complex (Figure 4a, left). Given the injected protein concentrations, we anticipated that REV7 might engage as a dimer, after which only one molecule is stably associated with SHLD3. Indeed, when using a dimerization deficient REV7^R124A^, we observed no discernible dissociation in the second phase (Figure 4a, right). These experiments highlight the extremely slow association and dissociation constants of the REV7-SHLD3^RBM^ interaction. We used the plateau values at the end of the dissociation phase to estimate the binding constant of the REV7-SHLD3^RBM^ interaction to be ~38 nM, in agreement with previous studies (Dai et al., 2020; Gupta et al., 2018) (Figure S3a). A stringent washing and regeneration protocol allowed for repeated experiments with increasing concentrations of REV7, which we used to calculate the association constant kon (Figure 4b,c). We observed no significant differences between the wild-type and dimerization mutant of REV7^R124A^, suggesting that dimerization itself does not influence the interaction kinetics of REV7-SHLD3^RBM^. This is reminiscent of the MAD2-mediated assembly of MCC where a dimerization mutant does not affect basal assembly (Piano et al., 2021). The binding constants determined for REV7-SHLD3^RBM^ were similar to the interaction kinetics of REV7 to REV3, suggesting that the interaction is not assisted by the client protein (Figure 4d,e) (Hara et al., 2010; Rizzo et al., 2018).

**Figure 4.**
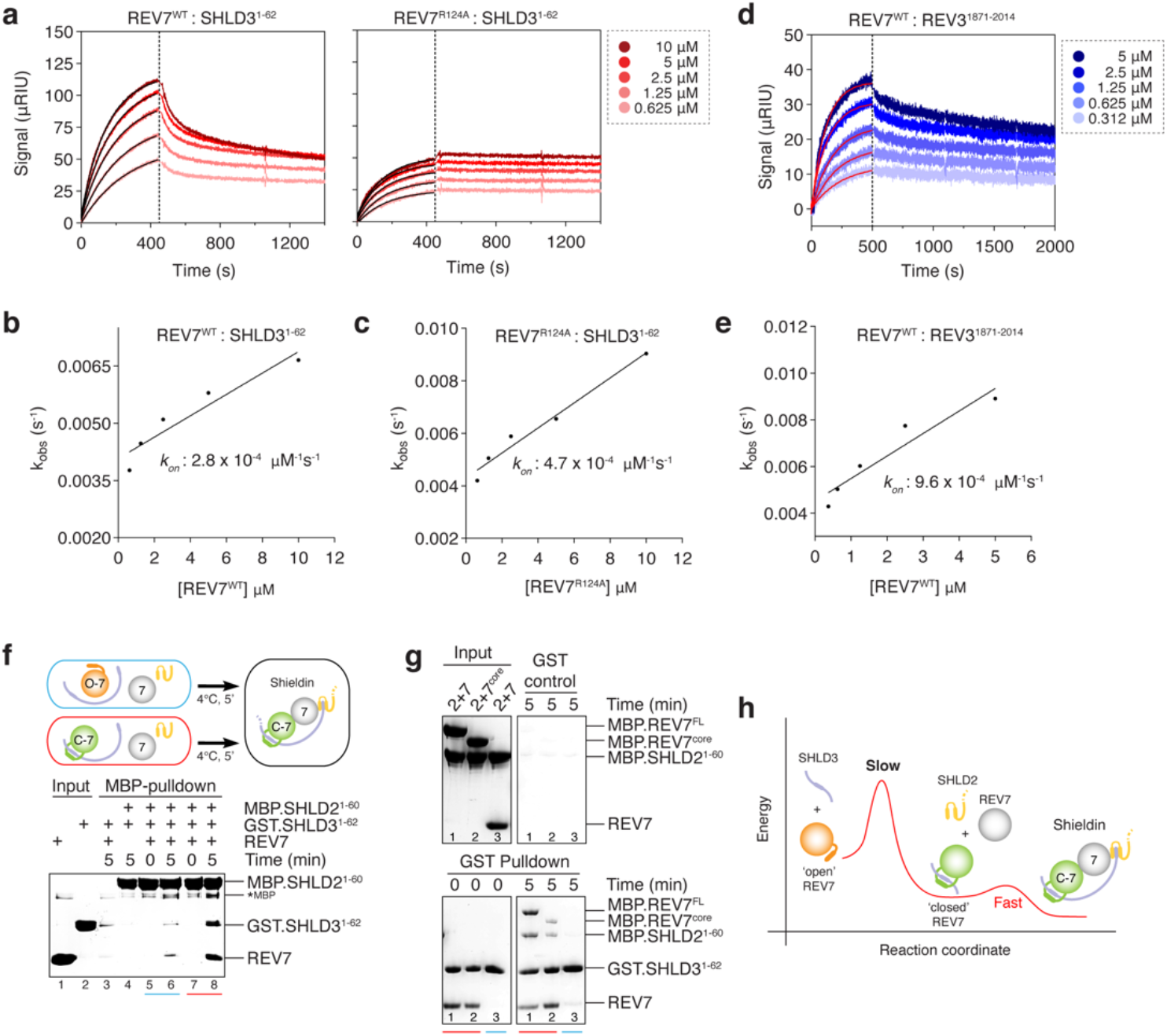
REV7-SHLD3 binding is rate limiting for Shieldin assembly. **a)** SPR sensograms obtained after injection of REV7^WT^ or REV7^R124A^ over immobilized SHLD3^1-62^ on a HC30M chip. **b)** Association kinetics of WT-REV7 with SHLD3^1-62^. Calculated from sensorgram in a). **c)** Association kinetics of R124A-REV7 with SHLD3^1-62^. Calculated from sensorgram in a). **d)** SPR sensorgram obtained after injection of REV7^WT^ over immobilized REV3^1871-2014^ on a HC30M chip. **e)** Association kinetics of WT-REV7 with REV3^1871-2014^. Calculated from sensorgram in d). **f)** MBP pulldown showing interaction kinetics between MBP-SHLD2^1-60^, REV7, GST-SHLD3^1-62^ or preformed GST-SHLD3^1-62^-REV7. **g)** GST pulldown showing interaction kinetics between preformed GST-SHLD3^1-62^-REV7, MBP-SHLD2^1-60^, and MBP-REV7^WT^ or REV7^core^. **h)** Schematic of the proposed kinetic model for Shieldin assembly. Shieldin assembly depends on slow conversion of ‘open’ REV7 to ‘closed’ REV7 on SHLD3 RBM followed by fast incorporation of SHLD2 through REV7 dimerization.

Next, we wondered if the incorporation of the second molecule of REV7 is equally slow. This second REV7 interaction to SHLD3 is mediated via SHLD2. Indeed, SHLD2 is not sufficient to interact strongly to SHLD3 by itself (Figure 4f, lane 4), so any interaction to SHLD3 is dependent on the incorporation of the second REV7 molecule. When allowing only a five-minute incubation, SHLD2 can indeed merely incorporate sub-stoichiometric amounts of REV7-SHLD3, due to the slow and incomplete REV7-SHLD3^RBM^ complex formation (Figure 4f, lane 6). To selectively monitor the second REV7 interaction, we pre-form a stoichiometric REV7-SHLD3^RBM^ complex in the absence of SHLD2, which selectively incorporates only the first REV7 molecule. When mixing the preformed complex with additional REV7 and SHLD2, we observed strongly increased amounts of SHLD3 and REV7 in a SHLD2 pulldown after a five-minute incubation, indicating that incorporating the second REV7 is much faster than the first (Figure 4f, compare lane 6 and 8). Moreover, incorporating the REV7^core^ construct, that lacks all mobile elements in REV7, as second the REV7 molecule does not affect the assembly kinetics of Shieldin (Figure 4g). This shows that in contrast to the first REV7 interaction, no conformer conversion and no mobile structural elements are necessary for the second REV7.

## Discussion

Shieldin is an important component of the DNA double-strand break repair machinery by modulating resection and thereby inducing repair via NHEJ. In this role, Shieldin affects the development and treatment of human disease, such as Fanconi Anemia and cancer, in which it determines the sensitivity of BRCA1-deficient cancers to treatment with PARP inhibitors (Noordermeer and van Attikum, 2019). Given these critical roles, understanding the requirements for proper Shieldin activity is important.

SHLD3 is the first Shieldin complex subunit to be recruited at DNA DSBs (Gupta et al., 2018). Here, we have shown that SHLD3 binds both single- and double strand DNA via a DNA binding domain at its C-terminus, which likely aids in the initial recruitment to double strand breaks. This initial promiscuous DNA binding event would provide flexibility for the Shieldin complex to allow the initial assembly on both extensively resected as well as poorly resected DNA ends. The downstream recruited Shieldin component SHLD2 brings in three putative tandem oligonucleotide/oligosaccharide-binding (OB) folds that specifically bind to single strand DNA, which likely aid to direct the Shieldin complex to the proper DSB site where it can inhibit resection mediated by EXO1 and DNA2/BLM nucleases.

SHLD3 is the receptor for the assembly of the Shieldin complex, with REV7 to be incorporated next (Gupta et al., 2018). We quantify the interaction kinetics between REV7 and SHLD3^RBM^, and show that this interaction is several orders of magnitude slower than typical protein-protein interactions (Schreiber et al., 2009). In surprising contrast to previous work, apo-REV7 in our purifications is exclusively in the ‘closed’ state and we have not been able to induce spontaneous conformational switching to the ‘open’ state (Clairmont et al., 2020). The ‘open’ REV7 state might be required as an intermediate before capture of the client proteins, therefore the ‘closed’ apo-REV7 could represent an auto-inhibited state which would further decrease assembly kinetics. AAA^+^-ATPase TRIP13 could function to overcome this putative auto-inhibited state and prime REV7 for incorporation (see below). Future work will be needed to study Shieldin assembly starting from different conformers in order to deconvolute its full assembly mechanism.

The measured association constants between REV7 and SHLD3^RBM^ are comparable to the interaction kinetics of related HORMA domain proteins, like mitotic protein MAD2 (Piano et al., 2021; Simonetta et al., 2009; Vink et al., 2006). REV7 captures SHLD3^RBM^ in a seat-belt conformation, which is similar to how MAD2 captures the CDC20 closure motif. The slow interaction between MAD2 and CDC20 is due to the significant structural remodelling needed for MAD2 to wrap its C-terminal tail around the interacting peptide motif. This reversible structural remodelling is the rate-limiting step that triggers the assembly of the stable but temporary signalling complex (the Mitotic Checkpoint Complex). External factors like MAD1, BUB1, MPS1 and TRIP13 can accelerate the structural conversion of MAD2, both at the assembly and the disassembly level, allowing dynamic control of signalling (Faesen et al., 2017; Kim et al., 2018; Lara-Gonzalez et al., 2021; Piano et al., 2021; Ye et al., 2015). We hypothesize that the same concept is conserved for REV7 in Shieldin, where the first interaction in the Shieldin complex, between REV7 and SHLD3^RBM^, needs to be established before Shieldin can self-assemble. Since the c-NHEJ pathway is completed just under 30 minutes in cells, this argues for the presence of as-of-yet unidentified accelerating factors of Shieldin assembly (Mao et al., 2008).

The transient and asymmetric dimerization between an ‘open’ and ‘closed’ MAD2 is an essential step in the mechanism to accelerate conformer conversion (Antoni et al., 2005; Faesen et al., 2017). Similar to MAD2, we have not observed significant changes in Shieldin assembly induced by the dimerization of REV7. Therefore, although dimerization is essential, it is not sufficient to accelerate conversion. However, in contrast to MAD2, Shieldin complex contains a stable REV7 dimer (de Krijger et al., 2021; Liang et al., 2020; Xie et al., 2021). Since this dimer could potentially create an additional rate-limiting step in Shieldin assembly, we were intrigued by the understanding the conformer identity and interaction mechanism of the second REV7. In the recent structure of SHLD2-SHLD3-REV7, the seatbelt of this second REV7 is only partly invisible, but the position of beta-sheet (residues 198-211) suggested that it too is in a closed conformation. We show that REV7 can form dimers between different conformer states, an ability not seen in MAD2 (Mapelli et al., 2007). Additionally, all these dimers could be incorporated into Shieldin, as long as one REV7 molecule could adopt the ‘closed’ seat-belt interaction. The first REV7 could juxtaposed by a second REV7 that does not require any of the mobile structural elements for incorporation in Shieldin, resulting in a fast incorporation. Any external factors that control the assembly of Shieldin, would therefore need to accelerate the conversion of the first REV7, and not the second.

AAA^+^-ATPase TRIP13 activity is needed for proper Shieldin function in cells (Clairmont et al., 2020; de Krijger et al., 2021). We have recently shown that the interaction of TRIP13 to Shieldin, requires a dimer of REV7 (de Krijger et al., 2021; Xie et al., 2021). TRIP13 is a conserved and generic HORMA remodelling factor that can open ‘closed’ HORMA domain proteins in an ATP-dependent manner (Alfieri et al., 2018; Ye et al., 2015). For example, the opening of closed MAD2 induces the disassembly of the MCC and ‘primes’ the HORMA domains for renewed capture of client proteins by adopting the ‘open’ state. We propose that only the remodelling of the REV7 bound to SHLD3^RBM^ would be necessary to induce Shieldin disassembly, as the interaction of the second REV7 is conformer-independent. The role of TRIP13 co-factor p31 in this mechanism is unclear, as it is not necessary for TRIP13 interaction in contrast to MAD2 (de Krijger et al., 2021). REV7 and p31 are both HORMA domain proteins, that are reported to weakly interact (Rizzo et al., 2018). Overall, this suggests an exchange mechanism, where p31 would replace the second REV7 in the first step of disassembling Shieldin, after which it would direct TRIP13 to open the remaining ‘closed’ REV7. Alternatively, but not mutually exclusively, TRIP13 could function to open ‘closed’ apo-REV7, which was thermodynamically the most stable conformer in our purifications.

Overall, these results provide insights in the early steps of Shieldin recruitment and assembly on DSBs. First-to-be-recruited SHDL3 likely uses its newly identified DNA binding domain as the platform to recruit the first molecule of REV7. The RBM of SHLD3 is captured by REV7 using a seat-belt interaction mechanism, which represents a rate-limiting step in the assembly of Shieldin. After overcoming this obligatory step, Shieldin self-assembles fast using a cooperative mechanism within REV7-SHLD2-3 (Figure 4h). Future work will be required to find factors that modulate the conversion of REV7 conformers in a timely fashion and to specifically regulate the assembly of its many client effector complexes in different pathways.

## Acknowledgements

We thank Chuna Ram Choudhary (University of Copenhagen) for kindly sharing plasmids containing the SHLD2 and SHLD3 genes and Andrea Musacchio (MPI-MP) for the biGBac DNA assembly plasmid toolkit. We thank Patrick Cramer (MPI-NAT) for access to the fluorescence plate reader and Äkta micro system and Faesen lab members for their helpful discussions. We thank Bastian Föhr for critically reading the manuscript. This work was supported by core funding from the Max Planck Society to A.C.F.

## Author contributions

V.S. and A.C.F conceived the project, V.S. performed all the experiments and prepared the figures under the supervision of A.C.F., V.S. and A.C.F. wrote the manuscript.

## Competing interests

Authors declare no competing interests.

## Material and Methods

### Expression of recombinant proteins and purification

All recombinant proteins used in this study were of human origin. REV7 mutants, REV3^1871-2014^, SHLD2 N-terminal, and SHLD3 N- and C-terminal constructs were expressed with an N-terminal hexahistidine-MBP or GST fusion-tag from pLIB at 16°C in *E.coli* LOBSTR strain (Andersen et al., 2013) for 16 hr after induction with 0.1 mM IPTG. Cells were lysed by sonication in buffer A containing 25 mM HEPES-NaOH (pH 7.5), 0.3 M NaCl, 0.5 mM TCEP (VWR lifesciences) and 1mM PMSF (Roche). After clearing, the lysate was loaded on a Hi-Trap metal chelating column (Cytiva). Bound proteins were eluted with an imidazole gradient. The MBP tag or GST tag was cleaved from protein constructs using PreScission protease overnight and subsequently separated using reverse-affinity purification. Protein containing fractions were pooled, and concentrated in 10 kDa MWCO concentrator (Merck) and loaded onto a Superdex-75 column (GE Healthcare) equilibrated with buffer B containing 10mM HEPES-NaOH pH 7.5, 0.15 M NaCl and 0.5 mM TCEP for size exclusion chromatography (SEC). Fractions containing purified REV7 and SHLD3 constructs were concentrated, flash-frozen and stored at −80°C until use. REV7 (both R124A and WT)-SHLD3-SHLD2^1-95^ complex was purified as GST fusion constructs from insect cell using biGBac expression system (Weissmann et al., 2016). Bacmid produced from DH10EMBacY cells was used to transfect Sf9 cells and produce baculovirus. Baculovirus was amplified through three rounds of amplification and used to infect Hi5 cells. Cells infected with the viruses were cultured for 72 h before harvesting. Purification of protein complexes was carried out using above protocol. Further polishing of the complex prior SEC and after tag cleavage with PreScission protease was carried out by loading onto a cation-exchange (CE) Resource-S column (GE Healthcare) equilibrated in 10mM HEPES (pH 7.5), 50 mM NaCl and 0.5 mM TCEP. Elution was carried out using 0.05-1 M NaCl gradient over 20 column volumes. Fractions containing purified REV7 (both R124A and WT)-SHLD3-SHLD2^1-95^ were concentrated, flash-frozen and stored at −80°C until use.

### *In-vitro* binding assays

For MBP-pulldown experiments, 1 μM MBP-SHLD3^45-60^ pre-adsorbed on Amylose beads was incubated for 30 minutes at 4°C with 2 μM of REV7, REV7^LL^, REV7^ΔN^ and REV7^ΔC^ in buffer B. After two washing steps of 0.5 ml each with buffer containing 10 mM HEPES (pH7.5), 0.5 M NaCl, 5% glycerol and 0.5 mM TCEP, complexes immobilized on beads were mixed with SDS gel loading buffer and analyzed by 12% SDS-PAGE. For GST-pulldown experiments, 1 μM GST-SHLD3^1-63^-REV7 pre-adsorbed on glutathione beads was incubated for 5 minutes at 4°C with 2 μM of MBP-REV7^core^or MBP-REV7 and 2 μM MBP-SHLD2^1-60^ in buffer B. After two washing steps of 0.5 ml each with buffer containing 10 mM HEPES (pH 7.5), 0.01% Tween 20, 0.5 M NaCl, 5% glycerol and 0.5 mM TCEP, complexes immobilized on beads were mixed with SDS gel loading buffer and analyzed by 12% SDS-PAGE.

### Static angle light scattering measurement

The SLS measurement was performed by coupling Superdex-200 column with VISCOTEK 305 TDA detector (Malvern). The run was performed in buffer B with prior calibration using BSA. The scattering was measured at 90° (right angle scattering) and 7° (low angle scattering) and data evaluation was carried out using OmniSEC v5.12 software (Malvern).

### Anion exchange chromatography

Purified REV7 mutants at 500 μg in total were subjected to anion-exchange (AE) chromatography using a 1 ml Hi-Trap Q column (Cytiva). Protein samples were resuspended in low salt buffer B prior loading onto Q column. Bound proteins were eluted using a 20-500 mM NaCl gradient over 20 column volumes. Fractions collected from the run were analyzed by 12% SDS-PAGE.

### Analytical size-exclusion chromatography

SEC runs were performed on Superdex-200 column equilibrated with buffer C (20mM Na-HEPES pH 7.5, 5% (v/v) glycerol, 100 mM NaCl and 0.5mM TCEP). Prior run, SHLD3-SHLD2^1-95^-REV7^R124A^ complex and 5,6-FAM labelled 50 bp ssDNA with the following sequence 5’-AAG GGG AGC GGG GGA GGA TAA TAG GAA GGG GAG CGG GGG AGG ATA ATA GG-3’ was incubated for 30 minutes on ice in dark. Fractions collected from SEC run were analyzed by 12% SDS-PAGE and scanned at 520 nm on Amersham Imager 680. For the REV7 dimerization experiments, 20 μM of REV7 or MBP-REV7 constructs were loaded on Superdex-200 3.2/300 column equilibrated with buffer B. Fractions collected on during SEC were analyzed by 12% SDS-PAGE. For Shieldin assembly experiments, 5 μM of MBP-SHLD2^1-60^, GST-SHLD3^1-62^, and REV7 mutants were incubated on ice for 60 minutes prior loading on the Superdex-200 3.2/300 column equilibrated with buffer B. Fractions collected during SEC run were analyzed by 12% SDS-PAGE. For Shieldin assembly using REV7^core^ construct, 5 μM of preformed REV7^ΔN^-SHLD3 was incubated with 10 μM of MBP-SHLD2^1-60^ and MBP-REV7^core^ on ice for 30 minutes prior loading on the Superdex-200 3.2/300 column equilibrated with buffer B. Fractions collected during SEC run were analyzed by 12% SDS-PAGE.

### Fluorescence anisotropy

A 5,6-FAM labelled ssDNA probe with the following sequence 5’-AGT GCC AGT GCC-3’ was purchased from Integrated DNA Technologies and was dissolved in deionized water. The dsDNA probe was generated by heating to 90°C and then slowly annealing equimolar amounts of 5’-GGC ACT GGC ACT-3’ with FAM labelled 5’-AGT GCC AGT GCC-3’. For binding affinity measurements, SHLD3-SHLD2^1-95^-REV7^R124A^/SHLD3^140-250^ was diluted in half-log steps in buffer C. Nucleic acids at a final concentration of 10 nM and SHLD3-SHLD2^1-95^-REV7^R124A^/SHLD3^140-250^ at a concentration range 0-56 μM were mixed on ice. The reaction was brought to a final volume of 25 μL and incubated in dark for 30 mins at room temperature. 18 μL of the reaction mixture was transferred to a Greiner 384 Flat bottom small volume plate. Fluorescence anisotropy was measured with an excitation wavelength of 470 ± 5 nm, an emission wavelength of 518 ± 5 nm, and a gain of 56. Each experiment was performed in triplicates and data was analysed using Graphpad Prism version 8. Binding curves were fit using a one site - specific binding equation. For anisotropy measurements, proteins at 20 μM were incubated with 10 nM of DNA probes. The reaction was brought to a final volume of 25 μL and incubated in dark for 30 mins at room temperature. The measurements were carried out as stated above. Statistical significance was performed using two-tailed student’s t test. Each experiment was performed at least in triplicates and data was analysed using Graphpad Prism version 8.

### Surface plasmon resonance measurements

Kinetic measurements were performed on a 2SPR Dual Channel system (XanTec bioanalytics GmbH). Purified SHLD3^1-62^ was immobilized on SCR HC30M chip using EDC-NHS coupling reaction. REV7^WT/R124A^ in two-fold dilutions was injected and analysed. Similar set-up was used for REV7-REV3 protein system with REV3^1871-2014^ immobilized on SCR HC30M chip. Buffer B supplemented with 0.05% Tween 20 (BIO-RAD) was used as a running buffer throughout SPR measurements. Data analysis was carried out using Trace Drawer software (XanTec bioanalytics GmbH).

### Statistics and reproducibility

Statistical analyses were performed with GraphPad Prism using the two-tailed student’s t-test. Significance; ns, not significant (*p*≥0.05), *\*p*<0.05, \**\*p*<0.01, \*\**\*p*< 0.001, *\*\*\*\*p*< 0.0001. Unless indicated otherwise, all gels are representative of at least two independent experiments, with uncropped gels shown in the Source data file.

---

**Figure S1.**
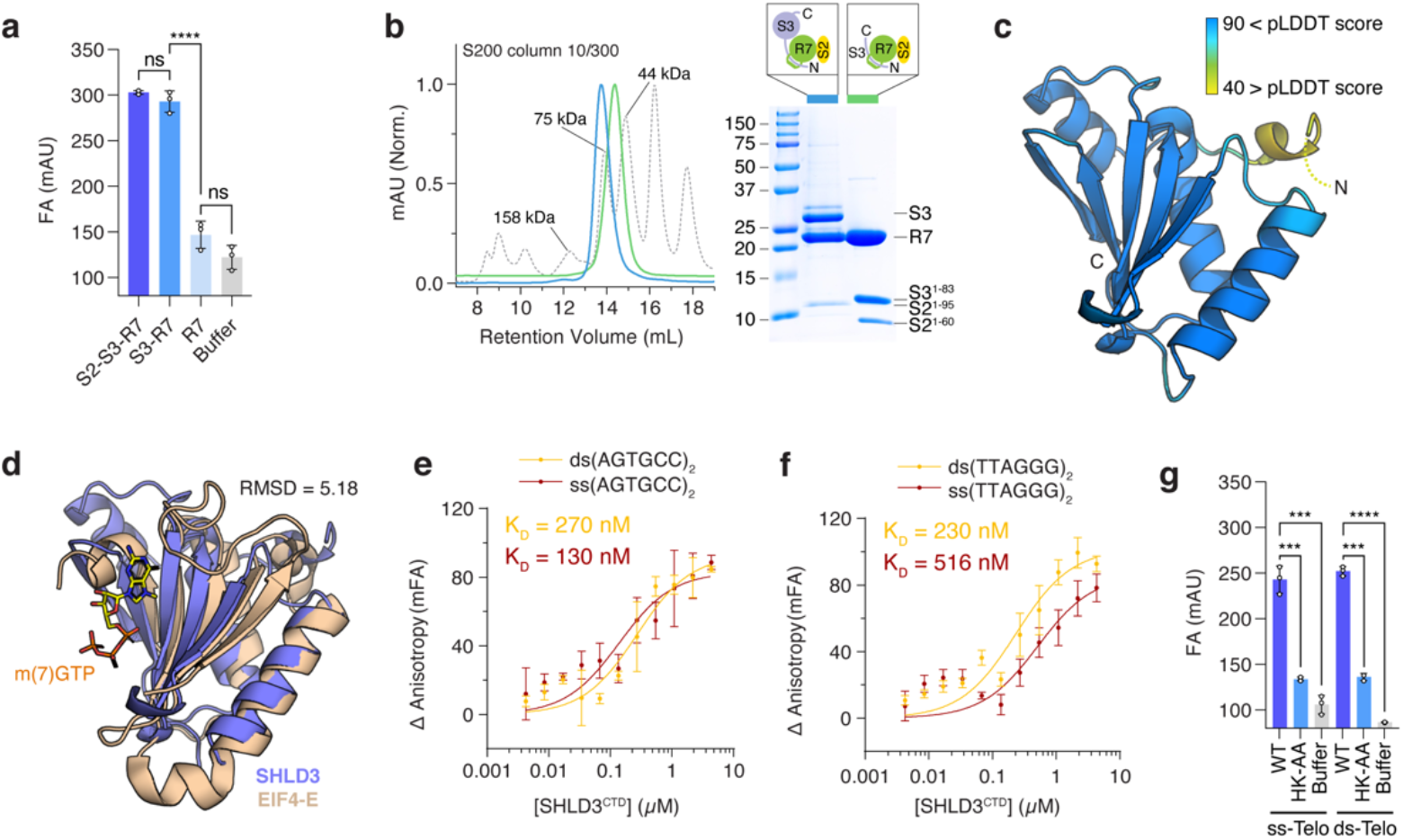
SHLD3 is a novel DNA binding protein, related to Figure 2. **a)** Fluorescence anisotrophy measurements of Shieldin complex, SHLD3-REV7, and/or REV7 for non-telomeric ssDNA substrates. Error bars represent s.d. (n = 3 independent experiments). Two-tailed Student’s test are indicated: ns, not significant, ****p<0.0001. **b)** Purification of stable SHLD3-SHLD2^1-95^-REV7 (blue curve) and SHLD3^1-83^-SHLD2^1-60^-REV7 (green curve) complex by size exclusion chromatography **c)** AlphaFold2 predicts a folded SHLD3 C-with a very high confidence (pLDDT > 90). **d)** Structural alignment of SHLD3^CTD^ and human EIF4-E (PDB: 5BXV) shows SHLD3^CTD^ adopts a fold similar to nucleotide binding translation factor. Proteins are coloured as follows SHLD3^CTD^ (marine blue), EIF4-E (beige), and m(7)GTP shown in stick representation. **e)** Fluorescence anisotrophy measurement of SHLD3^CTD^ for binding affinities for non-telomeric ss and dsDNA and **f)** for telomeric ss and dsDNA. Error bars represent s.d. (n = 3 independent experiments). **g)** Fluorescence anisotrophy measurements of SHLD3^CTD^ wildtype and mutants for telomeric ss- and dsDNA substrates. Error bars represent s.d. (n = 3 independent experiments). Two-tailed Student’s test are indicated: ***p<0.001, ****p<0.0001

**Figure S2.**
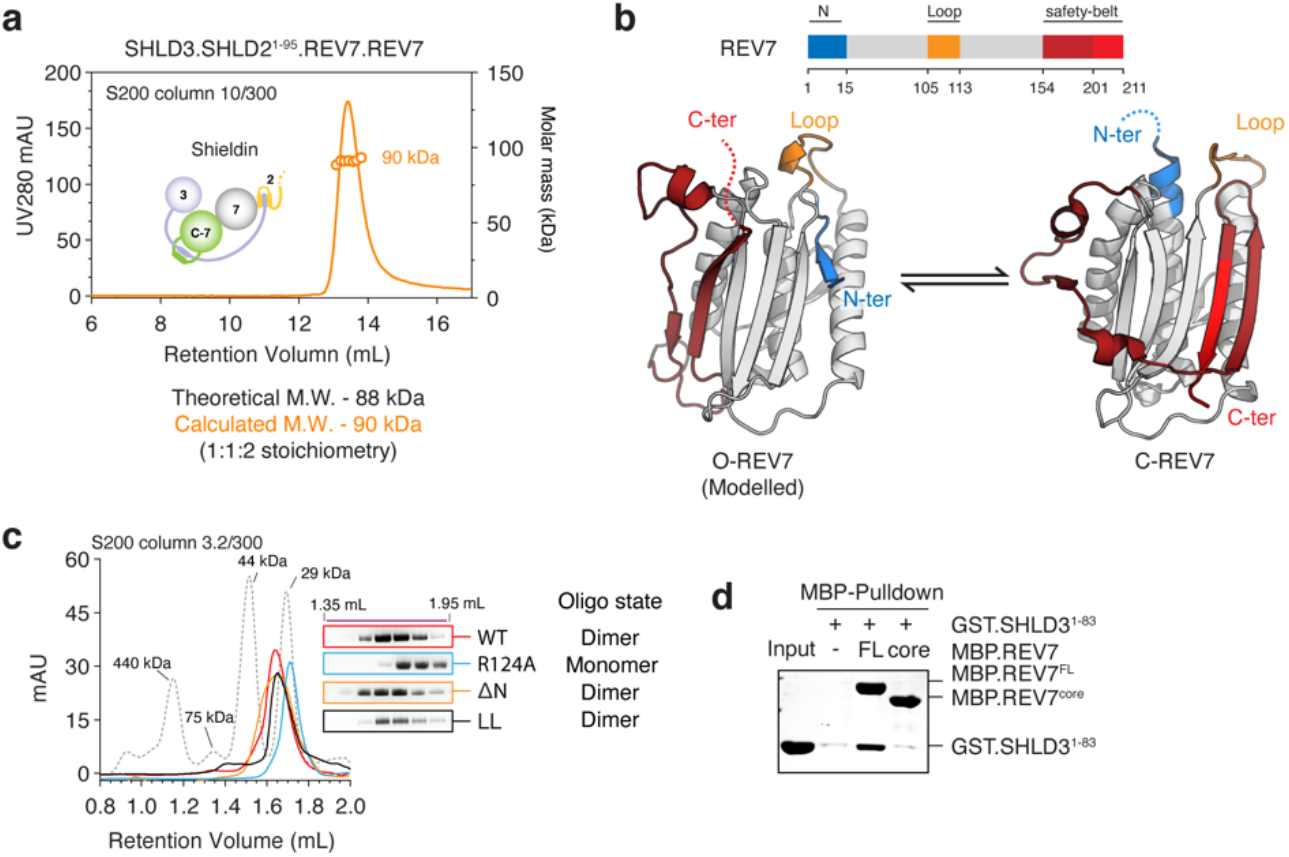
REV7 forms topology insensitive dimers, related to Figure 3. **a)** Measurement of absolute molar mass of SHLD3-SHLD2^1-95^-REV7 ternary complex using static angle light scattering coupled to size exclusion chromatography. Calculated molar mass of ~ 90 kDa predicts a dimer of REV7 present for each monomer of SHLD3 and SHLD2. **b)** Structural model of ‘open’ REV7 and crystal structure of ‘closed’ REV7. Conversion from ‘open’ to ‘closed’ conformer requires major structural rearrangement of N-terminal region (blue) and C-terminal safety belt region (red). Conversion requires the N-terminal region to thread through the loop region (orange). Inability to thread locks REV7 in ‘open’ state. **c)** Elution profile of REV7 protein mutants from size exclusion chromatography. The contents of consecutive 50 μL fractions eluting from 1.35 and 1.95 mL are shown. **d)** MBP pulldown showing the interaction between GST-SHLD3^1-83^ with MBP-REV7^WT/core^ mutants. REV7^core^ fails to bind GST-SHLD3^1-83^.

**Figure S3.**
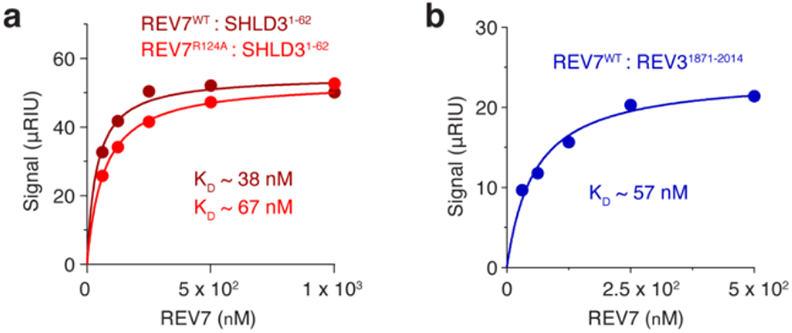
REV7 binds REV3 and SHLD3 with a slow on rate, related to Figure 4. **a)** Measurement of binding affinity of SHLD3^1-62^ to REV7^WT^ or REV7^R124A^. **b)** Measurement of binding affinity of REV3^1871-2014^ to REV7^WT^.

## Notes

### Competing Interest Statement

The authors have declared no competing interest.

## References

Alfieri C, Chang L, Barford D. 2018. Mechanism for remodelling of the cell cycle checkpoint protein MAD2 by the ATPase TRIP13. Nature 559:274-278–274–278. doi:10.1038/s41586-018-0281-1

Andersen KR, Leksa NC, Schwartz TU. 2013. Optimized E. coli expression strain LOBSTR eliminates common contaminants from His-tag purification. Proteins Struct Funct Bioinforma 81:1857-1861–1857–1861. doi:10.1002/prot.24364

Antoni AD, Pearson CG, Cimini D, Canman JC, Sala V, Nezi L, Mapelli M, Sironi L, Faretta M, Salmon ED, Musacchio A. 2005. The Mad1/Mad2 Complex as a Template for Mad2 Activation in the Spindle Assembly Checkpoint. Curr Biol 15:214-225–214–225. doi:10.1016/j.cub.2005.01.038

Ashkenazy H, Abadi S, Martz E, Chay O, Mayrose I, Pupko T, Ben-Tal N. 2016. ConSurf 2016: an improved methodology to estimate and visualize evolutionary conservation in macromolecules. Nucleic Acids Res 44:W344-W350–W344–W350. doi:10.1093/nar/gkw408

Clairmont CS, Sarangi P, Ponnienselvan K, Galli LD, Csete I, Moreau L, Adelmant G, Chowdhury D, Marto JA, D’Andrea AD. 2020. TRIP13 regulates DNA repair pathway choice through REV7 conformational change. Nat Cell Biol 22:87-96–87–96. doi:10.1038/s41556-019-0442-y

Dai Y, Zhang F, Wang L, Shan S, Gong Z, Zhou Z. 2020. Structural basis for shieldin complex subunit 3–mediated recruitment of the checkpoint protein REV7 during DNA double-strand break repair. J Biol Chem 295:250-262–250–262. doi:10.1074/jbc.ra119.011464

de Krijger I, Föhr B, Pérez SH, Vincendeau E, Serrat J, Thouin AM, Susvirkar V, Lescale C, Paniagua I, Hoekman L, Kaur S, Altelaar M, Deriano L, Faesen AC, Jacobs JJL. 2021. MAD2L2 dimerization and TRIP13 control shieldin activity in DNA repair. Nat Commun 12. doi:10.1038/s41467-021-25724-y

Dev H, Chiang T-WW, Lescale C, de Krijger I, Martin AG, Pilger D, Coates J, Sczaniecka-Clift M, Wei W, Ostermaier M, Herzog M, Lam J, Shea A, Demir M, Wu Q, Yang F, Fu B, Lai Z, Balmus G, Belotserkovskaya R, Serra V, O’Connor MJ, Bruna A, Beli P, Pellegrini L, Caldas C, Deriano L, Jacobs JJL, Galanty Y, Jackson SP. 2018. Shieldin complex promotes DNA end-joining and counters homologous recombination in BRCA1-null cells. Nat Cell Biol 20:954-965–954–965. doi:10.1038/s41556-018-0140-1

Erdős G, Pajkos M, Dosztányi Z. 2021. IUPred3: prediction of protein disorder enhanced with unambiguous experimental annotation and visualization of evolutionary conservation. Nucleic Acids Res 49:W297-W303–W297–W303. doi:10.1093/nar/gkab408

Faesen AC, Thanasoula M, Maffini S, Breit C, Müller F, van Gerwen S, Bange T, Musacchio A. 2017. Basis of catalytic assembly of the mitotic checkpoint complex. Nature 542:498-502–498–502. doi:10.1038/nature21384

Ghezraoui H, Oliveira C, Becker JR, Bilham K, Moralli D, Anzilotti C, Fischer R, Deobagkar-Lele M, Sanchiz-Calvo M, Fueyo-Marcos E, Bonham S, Kessler BM, Rottenberg S, Cornall RJ, Green CM, Chapman JR. 2018. 53BP1 cooperation with the REV7–shieldin complex underpins DNA structure-specific NHEJ. Nature 560:122-127–122–127. doi:10.1038/s41586-018-0362-1

Gu Y, Desai A, Corbett KD. 2022. Evolutionary Dynamics and Molecular Mechanisms of HORMA Domain Protein Signaling. Annu Rev Biochem 91:annurev-biochem-090920-103246. doi:10.1146/annurev-biochem-090920-103246

Gupta R, Somyajit K, Narita T, Maskey E, Stanlie A, Kremer M, Typas D, Lammers M, Mailand N, Nussenzweig A, Lukas J, Choudhary C. 2018. DNA Repair Network Analysis Reveals Shieldin as a Key Regulator of NHEJ and PARP Inhibitor Sensitivity. Cell 173:972-988.e23–972-988.e23. doi:10.1016/j.cell.2018.03.050

Hara K, Hashimoto H, Murakumo Y, Kobayashi S, Kogame T, Unzai S, Akashi S, Takeda S, Shimizu T, Sato M. 2010. Crystal Structure of Human REV7 in Complex with a Human REV3 Fragment and Structural Implication of the Interaction between DNA Polymerase ζ and REV1. J Biol Chem 285:12299-12307–12299–12307. doi:10.1074/jbc.m109.092403

Jumper J, Evans R, Pritzel A, Green T, Figurnov M, Ronneberger O, Tunyasuvunakool K, Bates R, Žídek A, Potapenko A, Bridgland A, Meyer C, Kohl SAA, Ballard AJ, Cowie A, Romera-Paredes B, Nikolov S, Jain R, Adler J, Back T, Petersen S, Reiman D, Clancy E, Zielinski M, Steinegger M, Pacholska M, Berghammer T, Bodenstein S, Silver D, Vinyals O, Senior AW, Kavukcuoglu K, Kohli P, Hassabis D. 2021. Highly accurate protein structure prediction with AlphaFold. Nature 596:583-589–583–589. doi:10.1038/s41586-021-03819-2

Kim DH, Han JS, Ly P, Ye Q, McMahon MA, Myung K, Corbett KD, Cleveland DW. 2018. TRIP13 and APC15 drive mitotic exit by turnover of interphase-and unattached kinetochore-produced MCC. Nat Commun 9:4354. doi:10.1038/s41467-018-06774-1

Lara-Gonzalez P, Kim T, Oegema K, Corbett K, Desai A. 2021. A tripartite mechanism catalyzes Mad2-Cdc20 assembly at unattached kinetochores. Science 371:64–67. doi:10.1126/science.abc1424

Liang L, Feng J, Zuo P, Yang J, Lu Y, Yin Y. 2020. Molecular basis for assembly of the shieldin complex and its implications for NHEJ. Nat Commun 11. doi:10.1038/s41467-020-15879-5

Luo X, Tang Z, Xia G, Wassmann K, Matsumoto T, Rizo J, Yu H. 2004. The Mad2 spindle checkpoint protein has two distinct natively folded states. Nat Struct Mol Biol 11:338–345. doi:10.1038/nsmb748

Malik R, Kopylov M, Gomez-Llorente Y, Jain R, Johnson RE, Prakash L, Prakash S, Ubarretxena-Belandia I, Aggarwal AK. 2020. Structure and mechanism of B-family DNA polymerase ζ specialized for translesion DNA synthesis. Nat Struct Mol Biol 27:913-924–913–924. doi:10.1038/s41594-020-0476-7

Mapelli M, Massimiliano L, Santaguida S, Musacchio A. 2007. The Mad2 Conformational Dimer: Structure and Implications for the Spindle Assembly Checkpoint. Cell 131:730-743–730–743. doi:10.1016/j.cell.2007.08.049

Mirdita M, Schütze K, Moriwaki Y, Heo L, Ovchinnikov S, Steinegger M. 2021. ColabFold - Making protein folding accessible to all. doi:10.1101/2021.08.15.456425

Musacchio A. 2015. The Molecular Biology of Spindle Assembly Checkpoint Signaling Dynamics. Curr Biol 25:R1002–R1018. doi:10.1016/j.cub.2015.08.051

Noordermeer SM, Adam S, Setiaputra D, Barazas M, Pettitt SJ, Ling AK, Olivieri M, Álvarez-Quilón A, Moatti N, Zimmermann M, Annunziato S, Krastev DB, Song F, Brandsma I, Frankum J, Brough R, Sherker A, Landry S, Szilard RK, Munro MM, McEwan A, de Rugy TG, Lin Z-Y, Hart T, Moffat J, Gingras A-C, Martin A, van Attikum H, Jonkers J, Lord CJ, Rottenberg S, Durocher D. 2018. The shieldin complex mediates 53BP1-dependent DNA repair. Nature 560:117-121–117–121. doi:10.1038/s41586-018-0340-7

Noordermeer SM, van Attikum H. 2019. PARP Inhibitor Resistance: A Tug-of-War in BRCA-Mutated Cells. Trends Cell Biol 29:820–834. doi:10.1016/j.tcb.2019.07.008

Piano V, Alex A, Stege P, Maffini S, Stoppiello GA, in ’t Veld PJH, Vetter IR, Musacchio A. 2021. CDC20 assists its catalytic incorporation in the mitotic checkpoint complex. Science 371:67-71–67–71. doi:10.1126/science.abc1152

Rizzo AA, Vassel F-M, Chatterjee N, D’Souza S, Li Y, Hao B, Hemann MT, Walker GC, Korzhnev DM. 2018. Rev7 dimerization is important for assembly and function of the Rev1/Polζ translesion synthesis complex. Proc Natl Acad Sci 115:E8191-E8200–E8191–E8200.

Schreiber G, Haran G, Zhou H-X. 2009. Fundamental Aspects of Protein-Protein Association Kinetics. Chem Rev 109:839–860. doi:10.1021/cr800373w

Setiaputra D, Durocher D. 2019. Shieldin – the protector of DNA ends. EMBO Rep 20. doi:10.15252/embr.201847560

Simonetta M, Manzoni R, Mosca R, Mapelli M, Massimiliano L, Vink M, Novak B, Musacchio A, Ciliberto A. 2009. The Influence of Catalysis on Mad2 Activation Dynamics. PLoS Biol 7:e1000010–e1000010. doi:10.1371/journal.pbio.1000010

Vink M, Simonetta M, Transidico P, Ferrari K, Mapelli M, Antoni AD, Massimiliano L, Ciliberto A, Faretta M, Salmon ED, Musacchio A. 2006. In Vitro FRAP Identifies the Minimal Requirements for Mad2 Kinetochore Dynamics. Curr Biol 16:755-766–755–766. doi:10.1016/j.cub.2006.03.057

Weissmann F, Petzold G, VanderLinden R, Brown NG, Lampert F, Westermann S, Stark H, Schulman BA, Peters J-M, others. 2016. biGBac enables rapid gene assembly for the expression of large multisubunit protein complexes. Proc Natl Acad Sci 113:E2564-E2569–E2564–E2569.

Xie W, Wang S, Wang J, de la Cruz MJ, Xu G, Scaltriti M, Patel DJ. 2021. Molecular mechanisms of assembly and TRIP13-mediated remodeling of the human Shieldin complex. Proc Natl Acad Sci 118:e2024512118–e2024512118. doi:10.1073/pnas.2024512118

Yang M, Li B, Liu C-J, Tomchick DR, Machius M, Rizo J, Yu H, Luo X. 2008. Insights into Mad2 Regulation in the Spindle Checkpoint Revealed by the Crystal Structure of the Symmetric Mad2 Dimer. PLoS Biol 6:e50. doi:10.1371/journal.pbio.0060050

Ye Q, Rosenberg SC, Moeller A, Speir JA, Su TY, Corbett KD. 2015. TRIP13 is a protein-remodeling AAA+ ATPase that catalyzes MAD2 conformation switching. eLife 4:e07367. doi:10.7554/eLife.07367

Mao Z, Bozzella M, Seluanov A, Gorbunova V. 2008. Comparison of nonhomologous end joining and homologous recombination in human cells. DNA Repair 7:1765-1771–1765–1771. doi:10.1016/j.dnarep.2008.06.018

